# Lipidomics of human adipose tissue reveals diversity between body areas

**DOI:** 10.1101/2020.01.20.912527

**Authors:** Naba Al-Sari, Tommi Suvitaival, Ismo Mattila, Ashfaq Ali, Linda Ahonen, Kajetan Trost, Trine Foged Henriksen, Flemming Pociot, Lars Ove Dragsted, Cristina Legido-Quigley

**Affiliations:** Steno Diabetes Center Copenhagen, Denmark; Dept. of Clinical Medicine, University of Copenhagen, Denmark; Capio CFR, Kgs. Lyngby, Denmark; Dept. Nutrition, Exercise and Sports, University of Copenhagen, Denmark; Institute of Pharmaceutical Science, King’s College London, UK

**Keywords:** Lipidomics, LC-MS, adipose tissue

## Abstract

**Background and aims:** Adipose tissue plays a pivotal role in storing excess fat and its composition reflects the history of person’s lifestyle and metabolic health. Broad profiling of lipids with mass spectrometry has potential for uncovering new knowledge on the pathology of obesity, metabolic syndrome, diabetes and other related conditions. Here, we developed a lipidomic method for analyzing human subcutaneous adipose biopsies. We applied the method to four body areas to understand the differences in lipid composition between these areas.

**Materials and methods:** Adipose tissue biopsies from 10 participants were analyzed using ultra-high-performance liquid chromatography coupled to quadrupole time-of-flight mass spectrometry. The method development included the optimization of the lipid extraction, the sample amount and the sample dilution factor to detect lipids in an appropriate concentration range. Lipidomic analyses were performed for adipose tissue collected from the abdomen, breast, thigh and lower back. Differences in lipid levels between tissues were visualized with heatmaps.

**Results:** Lipidomic analysis on human adipose biopsies lead to the identification of 187 lipids in 2 mg of sample. Technical variation of the lipid-class specific internal standards were below 5 %, thus indicating acceptable repeatability. Triacylglycerols were highly represented in the adipose tissue samples, and lipids from 13 lipid classes were identified. Long polyunsaturated triacylglycerols in higher levels in thigh (q<0.05), when compared with the abdomen, breast and lower back, indicating that the lipidome was area-specific.

**Conclusion:** The method presented here is suitable for the analysis of lipid profiles in 2 mg of adipose tissue. The amount of fat across the body is important for health but we argue that also the distribution and the particular profile of the lipidome may be relevant for metabolic outcomes. We suggest that the method presented in this paper could be useful for detecting such aberrations.

## 1. INTRODUCTION

Adipocytes are cells that form and store lipids in adipose tissue, they play a major role in energy homeostasis [1,2]. Obesity, which is characterized by increased storage of lipids in adipose tissue and modification of the metabolic functions of adipocytes, is a risk factor for several metabolic diseases, including atherosclerosis, cardiovascular ischemic disease, hypertension, hyperglycemia and insulin resistance in type 2 diabetes[1,3,4,5,6,7]. The molecular composition of adipose tissue may reveal its functionality and links to metabolic disease[8,2]. In addition, lipid deposition is important because individuals, who have more visceral adipose tissue, have a greater risk of metabolic disease[9]. Hence, global lipidomic analysis of multiple areas of adipose tissue may lead to a better understanding of metabolism in disease[10,11].

Lipidomics, together with genomics, transcriptomics, and proteomics, can give new biological insights and reveal novel metabolic pathways[12,13,14,15,16,17]. As an ‘-omics’ field, lipidomics is the approach of choice to understanding lipid biology[12,18,19,20]. Particularly, lipid composition of adipose tissue can reflect the long-term consumption of lipids[21]: adipose tissue has been shown to reflect dairy fat consumption at an annual level, whereas blood reflects the dietary intake over a time scale of weeks to months[22,23].

Lipidomics technologies can be used to measure hundreds of lipids in human biofluid or tissue[12,24]. However, few lipidomic analysis have been performed on human adipose tissue[25,26,27]. Lipidomics is a platform-dependent technique and numerous sample preparation methods have been reported[28,29,30,31,32]. Although there is no single technique to extract and cover the entire lipidome, the most common sample preparation techniques are variations of the liquid-liquid extraction (LLE) using the Folch extraction and the LLE using methyl *tert*-butyl ether (MTBE), originally introduced by Matyesh et al.[33,24]. Lipids from adipose tissue have also been extracted with a mixture of isooctane and ethyl acetate[29].

Here, we developed a method for a broad-coverage lipidomic analysis of human adipose biopsies that could be used at a clinical mass-spectrometry lab. The method was applied to adipose tissue samples from four body areas to identify possible compositional differences. The method detected nearly adipose tissue 200 lipids in this small cohort of obese participants. We observed that thigh subcutaneous adipose tissue had a different lipid profile compared to biopsies obtained from the back, abdomen and breast, whereas the three other areas were largely similar by their lipid composition.

## 2. MATERIALS AND METHODS

### 2.1 CHEMICALS

Methanol (MeOH) and water (H_2_O) of LC-MS grade was purchased from Honeywell (Morris Plains, NJ, USA). Chloroform (CHCI_3_), methyl-tert-butyl ether (MTBE) and Sodium chloride (NaCl) of reagent grade were purchased from Sigma-Aldrich (Steinheim, Germany).

Stock solutions (1 mg mL^−1^) of chosen lipid standards were prepared in CHCl_3_:MeOH (2:1, v/v). The following of these lipid standards were purchased from Sigma-Aldrich: 1,2-dimyristoyl-*sn*-glycero-3-phospho(choline-d13) (PC(14:0/d13)), 1,2,3-triheptadecanoylglycerol (TG(17:0/17:0/17:0)) and 3β-hydroxy-5-cholestene 3-linoleate (ChoE(18:2)). 1,2-diheptadecanoyl-*sn*-glycero-3-phosphoethanolamine (PE(17:0/17:0)), N-heptadecanoyl-D-*erythro*-sphingosylphosphorylcholine (SM(d18:1/17:0)),N-heptadecanoyl-D-*erythro*-sphingosine (Cer(d18:1/17:0)), 1,2-diheptadecanoyl-*sn*-glycero-3-phosphocholine (PC(17:0/17:0)), 1-heptadecanoyl-2-hydroxy-*sn*-glycero-3-phosphocholine (LPC(17:0)), 1-palmitoyl-d31-2-oleoyl-*sn*-glycero-3-phosphocholine (PC(16:0/d31/18:1)), 1-hexadecyl-2-(9Z-octadecenoyl)-sn-glycero-3-phosphocholine (PC(16:0e/18:1(9Z))), 1-(1Z-octadecenyl)-2-(9Z-octadecenoyl)-sn-glycero-3-phosphocholine (PC(18:0p/18:1(9Z))), 1-octadecanoyl-sn-glycero-3-phosphocholine (LPC(18:0)), 1-(1Z-octadecenyl)-2-do-cosahexaenoyl-*sn*-glycero-3-phosphocholine (PC(18:0p/22:6)). 1-stearoyl-2-linoleoyl-*sn*-glycerol (DG(18:0/20:4)) was purchased from Avanti Polar Lipids, Inc. (Alabaster, AL, USA) and tripalmitin-1,1,1-13C3 (TG(16:0/16:0/16:0)-13C3), trioctanoin-1,1,1-13C3 (TG(8:0/8:0/8:0)-13C3) and 1-palmitoyl-2-hydroxy-sn-Glycero-3phosphatidylcholine (LPC(16:0)) from Larodan AB (Solna, Sweden). A working standards solution (StdMix) was then prepared from these stock solutions into a concentration of 10 μg mL^−1^ in CHCl_3_:MeOH (2.1, v/v).

Calibration curves (at concentration levels of 100, 500, 1000, 1500, 2000 and 2500 ng mL^−1^) for the quantification of lipids were prepared using 12 different lipid standards. The following of these lipids were purchased from Sigma-Aldrich: 1,2,3-Triheptadecanoylglycerol (TG(17:0/17:0/17:0)) and 3β-Hydroxy-5-cholestene 3-linoleate (ChoE(18:2)), 3β-Hydroxy-5-cholestene 3-oleate (ChoE(18:1(9Z))), 5-Cholesten-3β-yl octadecenoate (ChoE(18:0)) and 5-Cholestene 3-palmitate (ChoE(16:0)). 1-Palmitoyl-2-Hydroxy-sn-Glycero-3-Phosphatidylcholine (LPC(16:0)) was purchased from Larodan and 1-hexadecyl-2-(9Z-octadecenoyl)-sn-glycero-3-phosphocholine (PC(16:0e/18:1(9Z))), 1-(1Z-octadecenyl)-2-(9Z-octadecenoyl)-sn-glycero-3-phosphocholine (PC(18:0p/18:1(9Z))), 1-octadecanoyl-sn-glycero-3-phosphocholine (LPC(18:0)), 1-(1Z-octadecenyl)-2-docosahexaenoyl-*sn*-glycero-3-phosphocholine (PC(18:0p/22:6)), 1-stearoyl-2-arachidonoyl-*sn*-glycero-3-phosphoinositol (PI(18:0/20:4)) and 1-stearoyl-2-linoleoyl-*sn*-glycerol (DG(18:0/18:2)) were purchased from Avanti Polar Lipids.

Additionally, the following quality control samples were prepared: blanks (solvent and matrix blanks), standard mixture samples, standard reference material (NIST 1950), pooled plasma samples and pooled adipose tissue samples. The pooled plasma samples and the NIST samples were prepared according to the following procedure: 10 μl aliquoted pooled plasma samples were thawed on ice. 10 μl 0.9% NaCl, 28 μl StdMix (10 μg/ml) were then added to the samples and extracted with 92 μl of CHCl_3_:MeOH (2:1, v/v).

### 2.2 SAMPLE PREPARATION AND SAMPLE ANALYSES

#### Ethical approval

Representative human adipose biopsy samples were collected anonymously from Capio CFR clinic in Lyngby during scheduled fat suctions requested by the donors, i.e. without registration of the donor identity, only donor age and site of fat suction. However, donors approved orally that waste materials may be used for methods development. Anonymous sampling means there is no way to trace the individual donor since no records or logs were made to associate donors and samples. When samples are completely anonymous and collected on the initiative of the donors they are not covered by the legislation on ethics and no ethical approval number was therefore obtained. Anonymous donors consented for waste material to be used for research.

Approximately 20g waste materials from the suctions were randomly collected. The 17 collected samples were from different body areas, abdomen (N= 6), breast (N=3), thigh (N=4) and lower back (N=4) from a total of N=10 participants. All the collected samples were transferred to Steno Diabetes Center Copenhagen (SDCC) on dry ice and were stored at −80 °C in collection tubes until analysis and the remaining samples were then discarded.

A section of adipose tissue sample was cut from each sample with a scalpel and loaded into the center of a tissue tube using a spatula. Transfer tubes were screwed onto the tissue tubes and the tubes were treated with liquid nitrogen for approximately 1 min to flash-freeze the tissue. The flash-frozen adipose tissue was cryopulverized using a CryoPREP™CP02 Pulverizer (Covaris Inc., USA), flash-frozen again and pulverized a second time. A given amount was weighed out from each sample and stored at −80°C until lipid extraction and analyses. All samples were randomized before sample preparation and again before analysis. Two different lipid extraction procedures (a modified Folch extraction procedure and a procedure based on extraction with MeOH:MTBE) were applied to the pulverized adipose tissue samples in order to determine the best procedure to use for further clinical studies.

The two different lipid extraction methods were initially performed on a sample from the abdomen of one subject using four replicates of cryopulverized samples (10 mg ± SD: 0.68 mg) in 1×, 5×, 10×, 20×, 40×, 80× and 100× dilutions. Four replicates of homogenate samples were extracted using the CHCl_3_:MeOH extraction procedure and four replicates of homogenate samples from one subject were extracted using the MeOH:MTBE extraction procedure. The modified Folch extraction procedure was performed as follows: first 200 μL of 0.9% NaCl was added to the cryopulverized adipose tissue samples. The samples were then spiked with 136 μL of a 10 μg mL^−1^ StdMix and extracted with 344 μL of CHCl_3_:MeOH (2:1, v/v). The samples were then vortex mixed, sonicated for 5 min, vortex mixed again and incubated on ice for 30 min. Finally, the samples were centrifuge at 10000 *rpm* for 3 min at 4 °C. The lower phase (100 μL) was then pipetted into a new Eppendorf tube and 100 μL of CHCl_3_:MeOH (2:1, v/v) was added. The samples were further diluted in a range of dilutions (1, 5, 10, 20, 40, 80 and 100 times) for different purposes as described below. The samples were stored at −80 °C until analysis.

The extraction procedure utilizing a mixture of MeOH and MTBE was somewhat different from the modified Folch extraction procedure. Here, 188 μL of H_2_O was added to the cryopulverized adipose tissue samples and the samples were then extracted with 975 μL of MeOH:MTBE:StdMix (225:750:35, v/v/v). The samples were vortex-mixed, sonicated for 5 min, vortex-mixed again and incubated on ice for 30 min. Finally, the samples were centrifuged at 10000 *rpm* for 3 min (4 °C) and 350 μL of the upper phase was pipetted into a new Eppendorf tube. The samples were then dried to completeness under nitrogen (N_2_). The samples were reconstituted in 110 μL of CHCl_3_:MeOH (2:1, v/v) and diluted serially (1, 5, 10, 20, 40, 80 and 100 times). The samples were stored at −80 °C until analysis. Once the samples had been prepared, they were analyzed using a previously published ultra-high-performance liquid chromatography quadrupole time-of-flight mass spectrometry method (UHPLC-Q-TOF-MS)[34,35,36,37]. All data were acquired using the MassHunter B.06.01 software by Agilent Technologies (Waldbronn).

### 2.3 DATA PROCESSING

The acquired data were pre-processed with MZmine (version 2.28) and peaks were annotated based on an in-house peak library and the LIPID MAPS^®^ database[38,39]. The peaks from the internal standards were detected by targeted analyses from the standard sample analyses to normalize each feature against internal standards in R. The other peaks in the samples were processed as follows: Mass detection was performed keeping the noise level at 1.5E3. Chromatogram builder used a minimum time span of 0.06 min, minimum height of 4.5E3, and *m/z* tolerance of 0.006 (or 10 ppm). Local minimum search algorithm was used for chromatogram deconvolution with a 70% chromatographic threshold, 0.05 min minimum RT range, minimum relative height 5%, minimum absolute height 7.5E3, minimum ratio of peak top/edge 1, and peak duration range 0.06-2.0 min.

Chromatograms were then deisotoped by using the isotopic peaks grouper algorithm with a *m/z* tolerance of 0.001 (or 2.0 ppm) and a RT tolerance of 0.05 min, following which the most abundant ion was kept. The peak list was filtered for exclude false signals and then row filtered for removing all rows which did not meet the requirement of minimum 1 peak in a row. Peak alignment was achieved using the join aligner method (*m/z* tolerance 0.006 (or 10.0 ppm), weight 2 for *m/z*, absolute RT tolerance 0.2 min), absolute RT tolerance of 1 min, and a threshold value of 1. Peak list row filtered again with minimum of 15 peaks in a row. The peak list was gap-filled with the same RT and *m/z* range gap filler (*m/z* tolerance at 0.006 or 10.0 ppm). Peak list was filtered again, and row filtered. Finally, the peak list was annotated using internal library with an m/z tolerance of 0.006 m/z or 10.0 ppm and a RT tolerance of 0.2 min.

The peak list was imported to R and each feature was normalized against an internal standard. The annotation included features, which had the same RT and m/z value as compounds in NIST 1950 serum samples^40^. Features that corresponded to equivalent standards injected during the sequence were labeled as Level 1 and features with structure information were labeled as Level 2^41^. Coefficient of variation (or, relative standard deviation; %RSD) for peak areas and retention times of lipid-class specific internal standards were calculated.

### 2.3 STATISTICAL ANALYSIS

Statistical analysis were performed with R (http://www.r-project.org/)[42]. The overall variation in samples were visualized in a principal component analysis (PCA) plot with annotation of the body area using the FactoMineR package[43]. Lipid-specific variation between body areas was assessed with lipid-wise linear regression models using the limma package, and the resulting model coefficients and their statistical significance were visualized on heatmaps using the ggplot2 package[44,45]. Results of the statistical tests were corrected for multiple testing using the Benjamini-Hochberg method.

## 3. RESULTS AND DISCUSSION

### 3.1 METHOD DEVELOPMENT

There are several protocols for lipidomics analyses suitable for the isolation and purification of lipids from blood and tissue samples[24,34,46,47,48]. The most widely used are based on the Folch extraction and an MTBE based method described by Matyash *et al*.[*24,49*]. To the knowledge of the authors, neither of these methods have so far been applied to human adipose tissue from different body sampling sites.

The aim of this study was to establish an optimal analytical procedure for multi-area adipose tissue lipidomics. Initially, two different extraction techniques (a modified Folch technique and an MTBE based technique) and two sample amounts and serial sample dilutions were tested. The optimization was made considering the number of annotated lipid features (unique retention time and *m/z* pairs) while minimizing peak saturation.

Both extraction methods - the CHCI_3_:MeOH and the MeOH:MTBE - consistently gave the same features. The results also suggested that 10 mg resulted in a saturation of peaks. A 20× dilution was deemed as the best dilution when considering the lipid features and lipid class abundance. The CHCl_3_:MeOH method with a sample amount of 2 mg and a dilution factor of 20× was hence determined as the optimal approach. In this protocol we aimed to minimize the starting material and focus on lipids that are abundant in adipose tissue, such as the triacylglycerols.

### 3.2 OVERALL LIPID COMPOSITION IN ADIPOSE TISSUE

Adipose tissue samples from different body areas were analyzed using the optimized method. The areas (abdomen, *N* = 6; breast, *N* = 3; lower back, *N* = 4; and thigh, *N* = 4) resulted in 187 lipid features from a 2 mg sample. These lipid features belonged to 13 major classes of lipids dominated by the triacylglycerols (TGs; S2 Tables 1 and 2). Also the following lipid classes were detected: diacylglycerols (DGs), ceramides (Cers), fatty acids (FAs), hexocyl-ceramides (HxCers), lyso-phosphatidylcholines (LPCs), phosphatidylcholines (PCs), alkyl-acyl phosphatidylcholines (PC-Os or PC-Ps), phosphatidylglycerols (PGs), phosphatidylethanolamines (PEs), alkyl-acyl phosphatidylethanolamines (PE-Os or PE-Ps), phosphatidylserines (PSs) and sphingomyelins (SMs).

Data from positive ion mode, which was analyzed further, covered eight of these lipid classes (DGs, HxCers, LPCs, PCs, PC-Os or PCPs, SMs and TGs). The relative numbers of identified lipids from these classes are shown in Fig 1 and all the identified lipids from positive as well as negative ion mode are listed in S2Tables 1 and 2. Analysis of technical variation indicated that the method was reproducible throughout the sample set. The coefficient of variation (or, relative standard deviation; RSD; %) of the peak areas were on average 4.8% for the internal standards (Table 1; 2.9% for the adipose tissue pooled samples and 6.7% for standard plasma samples; NIST).

**Fig 1:**
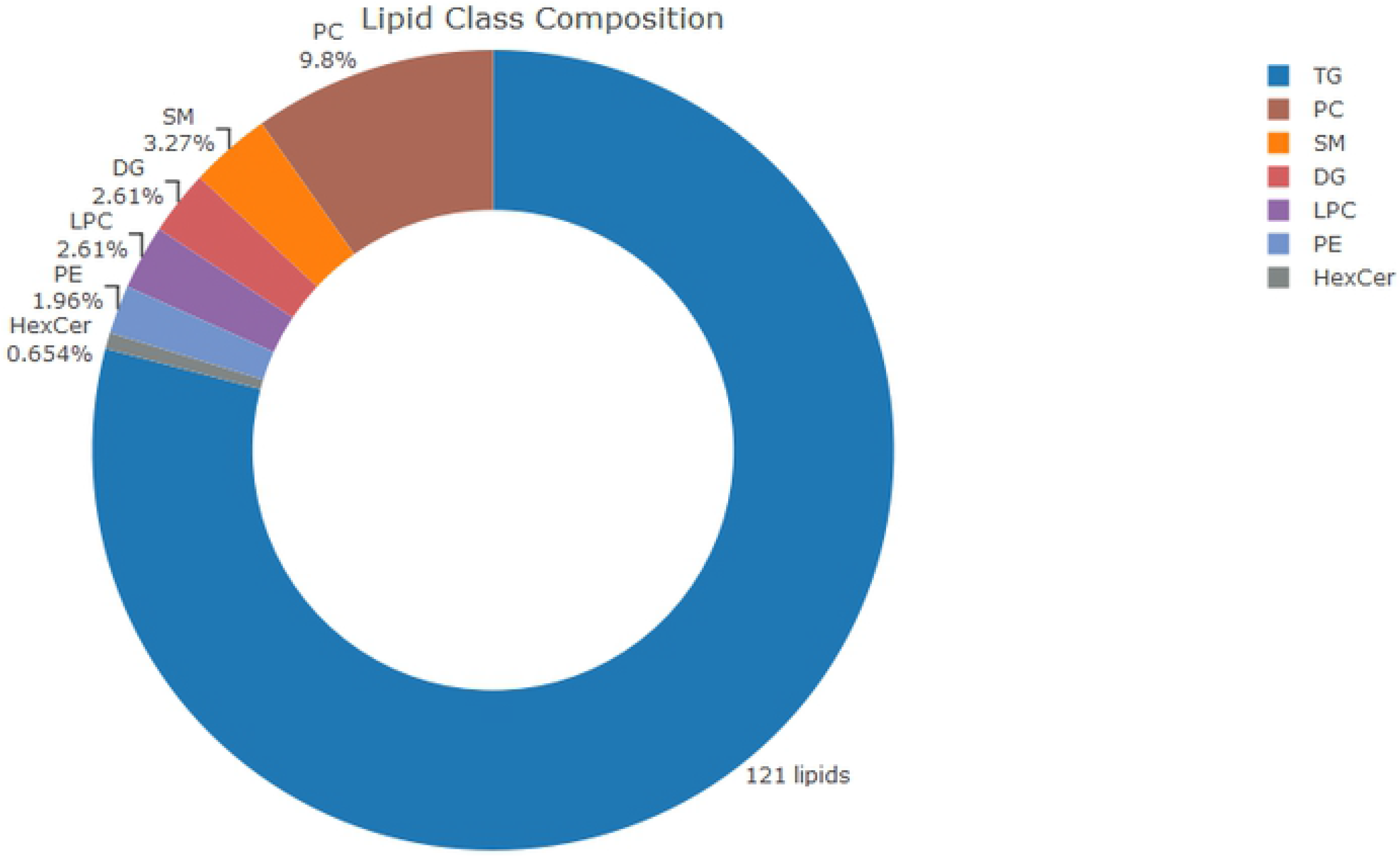
Donut plot showing numbers of lipids detected from each class and their proportion (%) among the identified lipids in adipose tissue. The result is from the optimized method.

A principal component analyses (PCA) of the lipid profiles revealed a multivariate pattern of the overall variation in the data (Fig 2). The results show that the adipose biopsy samples from the thigh clustered in the lower half of the projection and the pooled adipose tissue samples (crosses) clustered in the middle as an indication of method reproducibility.

**Fig 2:**
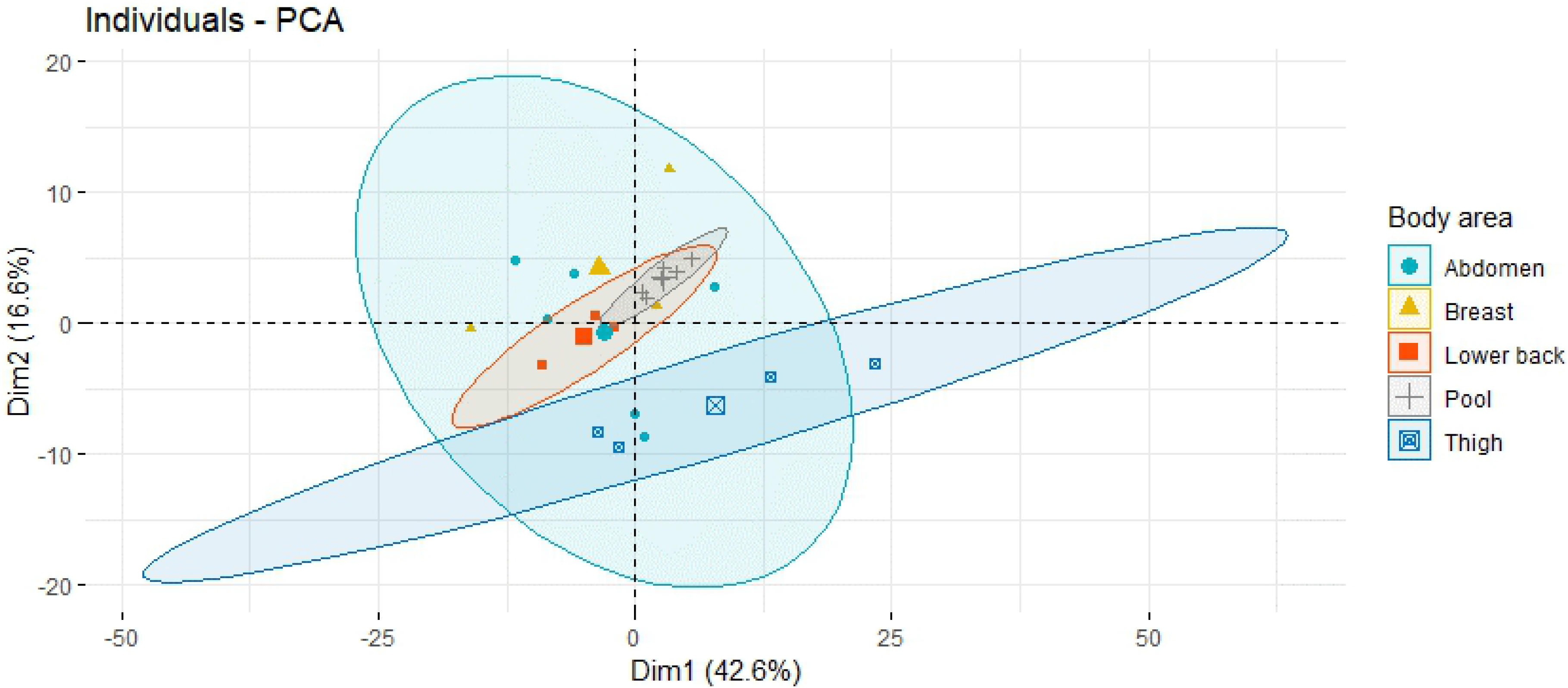
Principal component analysis (PCA) plot showing individual and technical variation (pooled samples as grey crosses). PCA summarizes the multivariate pattern of overall variation in the data that come from adipose tissue samples (n=17) from four body areas. The first two principal axes are shown on the x-axis and y-axis, respectively. Thigh (blue squares) is distinct from abdomen (blue circles), breast (yellow triangles) and lower back (red squares). The centroids of the sample groups are shown as larger-sized symbols and the two-standard-deviation (95 %) confidence intervals of the sample groups around the centroids are shown as ellipsoids.

Next, lipid levels were individually compared between the body areas to find out, whether the body’s lipid composition is location-specific. Levels of each lipid species were compared pair-wise between different areas. These comparisons are illustrated in the diagram of Fig 3A.

**Fig 3.**
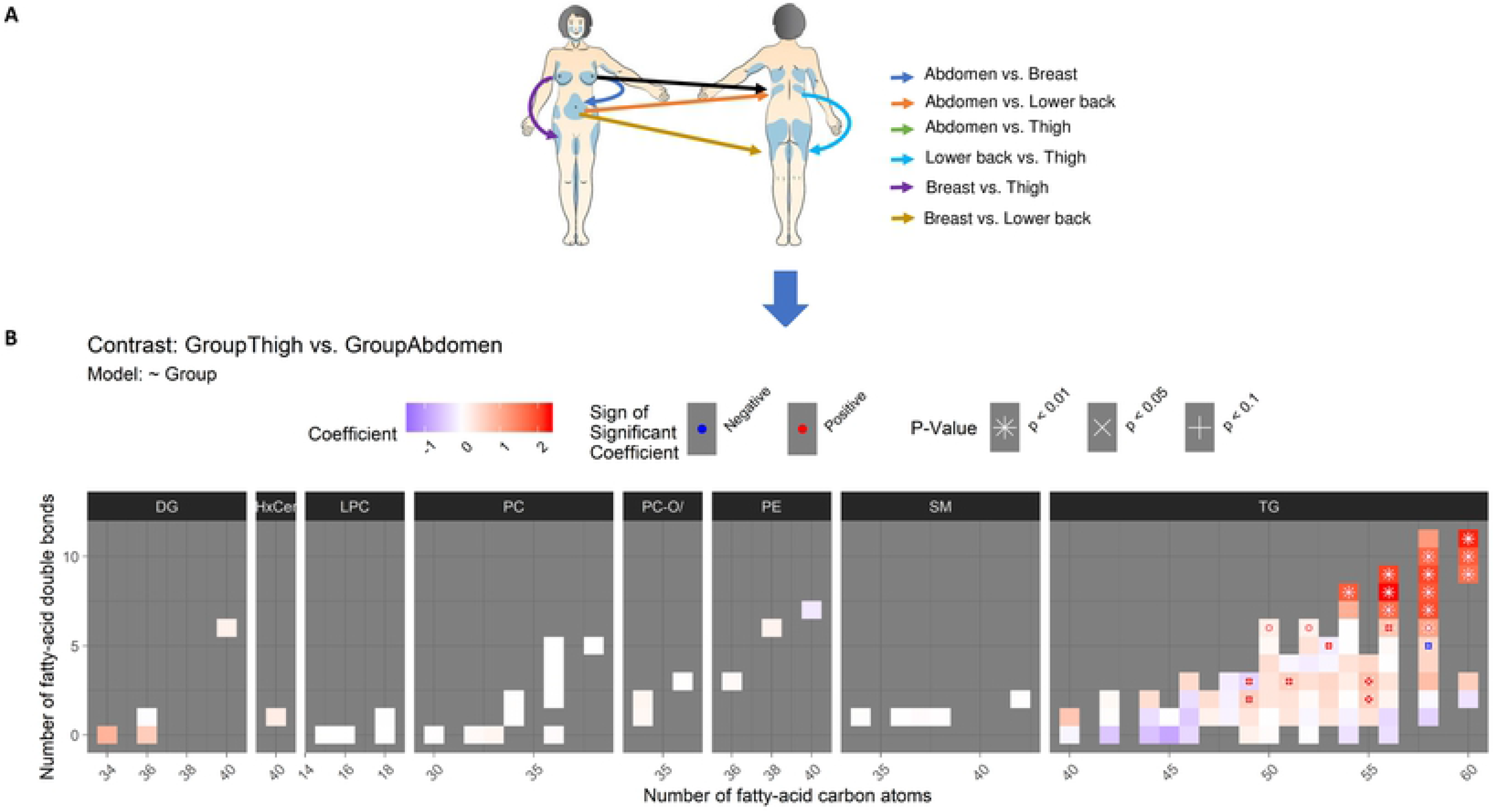
(A) Diagram illustrating all the pair-wise comparisons between adipose tissue samples from different body areas. Results of these comparisons are shown in Fig 3B, 4 and S1 Fig 1. (B) Difference in the lipidome between thigh and abdomen. Lipids are grouped by their class into panels (in columns from DG to TG). Within each panel, each colored rectangle corresponds to one lipid species, and its location in the x-axis and y-axis, respectively, shows its size (number of carbon atoms in the fatty acid chains) and its level of unsaturation (number of double bonds). Blue, red and white rectangles, respectively, indicate lower, higher and same levels in thigh compared to abdomen (based on the regression coefficient from the linear regression model with the body area as an independent variable and the lipid level as the dependent variable. Statistical significance of the difference is annotated by the symbols “*,” “x” and “+,” respectively, corresponding to p < 0.01, 0.05 and 0.1. For instance, the triacylglycerol TG(60:11), located in the x-and y-coordinates 60 and 11, respectively, (the top-rightmost corner) in the TG panel, has a total of 60 carbon atoms and 11 double bonds (i.e., unsaturated bonds) in its fatty-acid chains. The lipid TG(60:11) has a clearly higher level in thigh compared to abdomen (red color of the rectangle) with a statistical significance of p < 0.01 (annotation with the character “*”).

Thigh appeared the most distinct adipose tissue among the body areas. Thereby, we start by describing the distinct lipid profile of the adipose tissue in thigh.

### 3.3 DIFFERENCES BETWEEN THIGH AND OTHER BODY AREAS

First, thigh was compared to abdomen (Fig3B). The levels of all long polyunsaturated triacylglycerols (TGs) with more than 55 carbon atoms and five double-bonds were higher in thigh, and in all but two of these 13 TGs, the difference was significant at p < 0.01 after correction for multiple testing. This pattern can be seen in the top-right corner of the TG panel in Fig 3B with positive model coefficients (in red color) of these TG lipids. Furthermore, five mid-sized polyunsaturated TGs with 50 to 55 carbon atoms and two to six double bonds had higher levels in thigh at p < 0.05 (see the red-colored middle-right part of the TG panel in Fig 3B).

Second, the difference between thigh and lower back adipose tissue was broadly similar to the difference between thigh and abdomen (Fig 4A). Also when compared to lower back, thigh had higher levels of long polyunsaturated TGs (see the top-right part of the TG panel in Fig 4A). Although three of these TGs were higher only at p < 0.05, nine other TGs were higher at p < 0.01. More mid-sized polyunsaturated TGs (with 50 to 55 carbon atoms and two to six double bonds) were different between thigh and lower back than thigh and abdomen: The levels of nine mid-sized polyunsaturated TGs (with 50 to 55 carbon atoms and two to six double bonds) were higher in thigh at p < 0.05 (see the middle-right part of the TG panel in Fig 4A). This was more than the six, which were previously found different between thigh and abdomen. Unlike against abdomen, TG(58:5) was lower in thigh compared to lower back (p < 0.01; see the TG panel of Fig 4A at x-axis value 58 and y-axis value 5).

**Fig 4.**
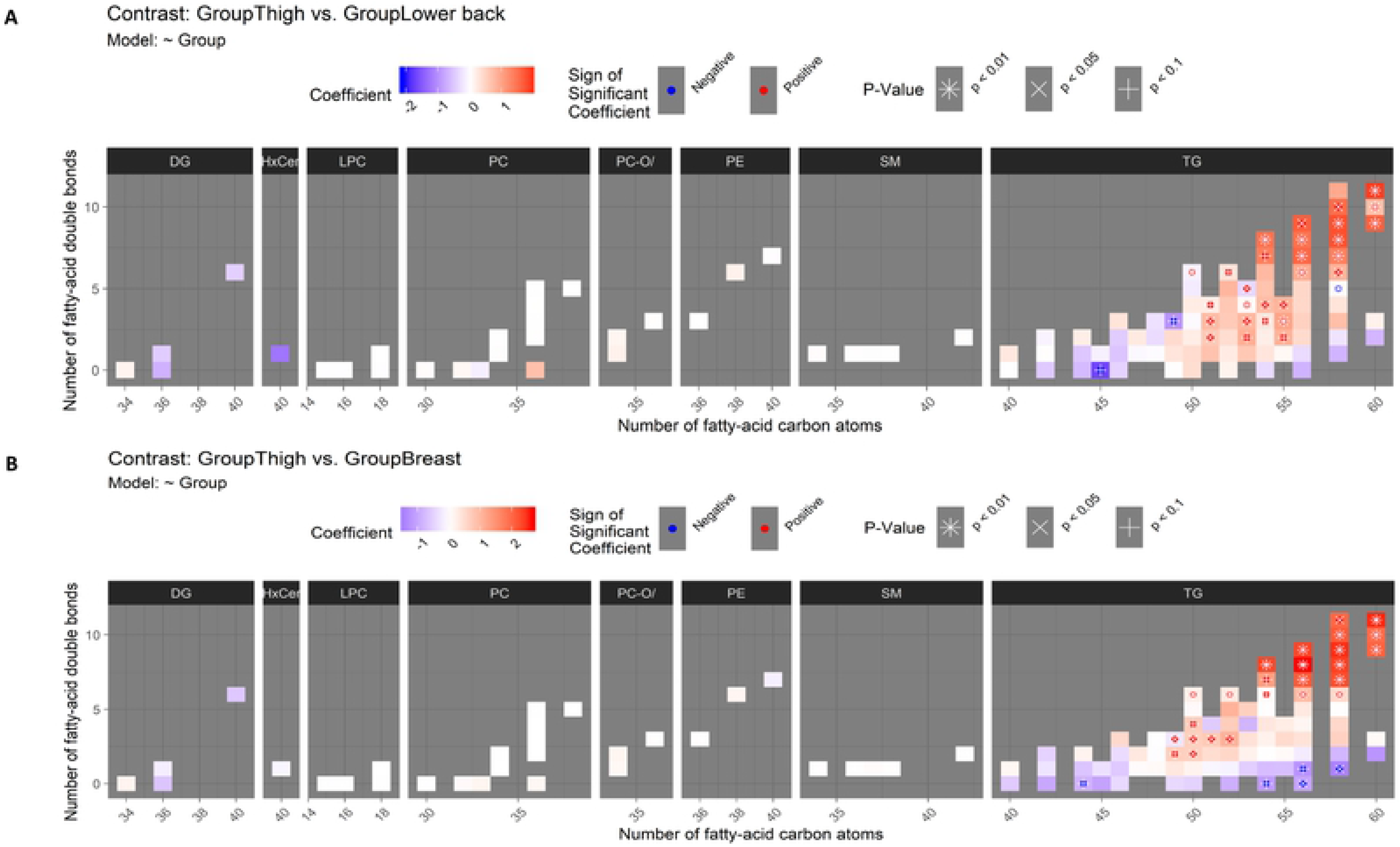
(A) Difference in the lipidome between thigh and lower back. See instructions for interpreting the figure in the caption of Fig 3B. (B) Difference in the lipidome between thigh and breast.

Again, in the comparison between thigh and breast, long polyunsaturated TGs appeared with clearly higher levels in thigh (Fig 4B): all 13 of these TGs were higher at p < 0.05 and only the weakest difference with TG(58:11) did not reach p < 0.01. Also a group of mid-sized TGs was again at higher levels in thigh than in breast but in this comparison the difference was confined to TGs with 49 to 52 carbon atoms instead of the up to 55 carbon atoms observed in the previous comparisons. The lower level of saturated and monounsaturated TGs in thigh compared to breast was more pronounced than in the comparisons involving thigh (see the bottom-part of the TG panel in Fig 4B): TG(56:0) and TG(58:1) were lower in thigh at p < 0.05. Other lipid classes did not have differences between thigh and other areas at p < 0.05 although non-significant patterns in DGs, HexCers and PCs were visible in Fig 3B and 4.

### 3.5 DIFFERENCES BETWEEN LOWER BACK, BREAST AND ABDOMEN

Only minor differences were observed between lower back and breast, when putting to the perspective of the rather rich signature of differences between thigh and other body areas. At p < 0.05, lower back had higher levels of the polyunsaturated TGs 52:6, 56:7 and 58:8 (see the top-right quarter of the panel in the “Lower Back vs. Breast” column and “TG” row in the S1 Fig 1). In a broader pattern of non-significant differences, lower back had generally higher levels of long polyunsaturated TGs and lower levels of long and medium-sized saturated and low-unsaturated TGs (see the right side of the panel in the “Lower Back vs. Breast” column and the “TG” row in the S1 Fig1).

Finally, no differences between lower back and abdomen or breast and abdomen were statistically significant (see the “Lower Back vs. Abdomen” and “Breast vs. Abdomen” columns in S1 Fig 1).

### 3.6 SUMMARY OF DIFFERENCES BETWEEN BODY AREAS

In summary, major differences in the lipid composition were observed between thigh and other body areas. The levels of long polyunsaturated TGs were consistently higher in thigh than in other areas and higher levels were also observed in medium-sized polyunsaturated TGs. In contrast, a group of saturated and monounsaturated TGs were lower in thigh than in the other body areas. This observation is noteworthy as Karastergiou *et al*. have shown that individuals with fat accumulated to the thighs show a lower risk for metabolic disease when compared to the accumulation of abdominal fat [50].

The current study was initiated to optimize the methodology for multi-area adipose tissue analysis and to explore, whether different body areas are comparable for adipose tissue collection. One strength of this study is that we had sufficient human adipose tissue from a single subject to optimize the methodology on homogenous material. Moreover, we tested both the major extraction methods with several body areas to assess potential area-specific variation.

A weakness of the study is its size and the fact that subjects were not a defined cohort; as a methodology development study it was designed with a small number of volunteers. The participants had adipose tissue removed for various reasons but the majority received a cosmetic operation to reduce excess adipose tissue in the central body, thighs or breasts. Hence, the findings of differences in TGs across the body areas should be studied further and their relationship to metabolic disease should be investigated. It would also be desirable to include a sufficient number of participants with simultaneous fat suction one of multiple body areas to enable the within-person comparison between body areas for improved statistical power.

## 4. CONCLUSIONS

This study showed that 2 mg of adipose tissue can produce good quality lipidomics data, acquiring relative levels of almost 200 lipid species belonging to eight lipid classes. The results suggest that adipose tissue in the thigh has a distinct composition of lipids with a higher amount of long polyunsaturated triacylglycerols (TGs), which is different from the lipid composition stored in the body’s trunk.

Further, a group of saturated and monounsaturated TGs were in lower levels in thigh compared to breast. This trend was also visible in other comparisons against thigh, although not significant, indicating generally somewhat lower levels of saturated and monounsaturated TGs in thigh. This observation is noteworthy as Karastergiou *et al*. have shown that individuals with fat accumulated in the thigh have a lower risk for metabolic disease when compared to abdominal accumulation of fat [50]. With the limited number of participants, though, it was not possible to investigate this and other patterns of individual variation further with the present study.

We hypothesize that aberrations in the balance of lipid species across the body could hold a key to further understanding the complications of metabolic syndrome. Not only the absolute amount of specific lipids in the body but also their distribution in the adipose tissue across the body might have implications on metabolic outcomes.

As routine clinical assessment of biomarkers from omics technologies becomes increasingly commonplace, we argue that the next frontier of personalized medicine will be to gain a deeper understanding of the balance of biomarkers across the active areas of the body. Depending on the condition, this could be between the blood and tissue, between different organs, or as in this work, between different areas of the same tissue type.

For metabolic disease, the distribution and location-specific content of adipose tissue could explain some missing links to the development of co-morbidities, such as cardiovascular disease, non-alcoholic liver disease and type 2 diabetes. In addition to adipose tissue, also the fat composition of the liver tissue has been shown to play an important role in the development of the metabolic co-morbidities [51]. However, biopsy sampling from the liver is significantly more intrusive an operation than that from the adipose tissue, and thereby it does not carry similar potential for broader adaption in the clinical practice.

On the other hand, increasing numbers of persons with obesity go through liposuction and bariatric operations [52], giving an opportunity also for the routine collection of adipose tissue material for profiling purposes. With the expense and speed of omics profiling technologies becoming increasingly applicable for clinical research and its translation to clinical practice, we expect that the hidden information also from the adipose tissue could be unleashed for the benefit of every patient. The ideas presented here are warranted for further investigation – now possible with the methodology reported in this paper.

